# Diversity of *Treponema denticola* and other oral treponeme lineages in subjects with periodontitis and gingivitis

**DOI:** 10.1101/2021.06.24.449858

**Authors:** Huihui Zeng, Yuki Chan, Wenling Gao, W. Keung Leung, Rory M. Watt

## Abstract

More than 75 species/species-level phylotypes of taxa belonging to the genus *Treponema* inhabit the human oral cavity. *Treponema denticola* is commonly associated with periodontal disease, but the etiological roles and ecological distributions of other oral treponemes remain more obscure. Here, we compared the clinical distributions of phylogroup 1 and 2 oral treponemes in subgingival plaque sampled from Chinese subjects with periodontitis (n=10) and gingivitis (n=8), via the sequence analysis of the highly-conserved *pyrH* ‘housekeeping’ gene. Two PCR primer sets that respectively targeted oral phylogroup 1 and 2 treponeme *pyrH* genes were used to construct plasmid clone-amplicon libraries for each subject, which were sequenced for bioinformatic analysis. 1,204 quality-filtered, full-length *pyrH* gene sequences were obtained from the cohort (median: 61.5; range: 59-83 of cloned *pyrH* sequences per subject), which were assigned to 34 ‘*pyrH* genotypes’ (pyrH001-pyrH034; 97% sequence-identity cut-off). 18 *pyrH* genotypes (536 *pyrH* sequences) corresponded to phylogroup 1 treponeme taxa (including *Treponema vincentii, Treponema medium*). 16 pyrH genotypes (668 *pyrH* sequences) corresponded to *T. denticola* and other phylogroup 2 treponemes. Periodontitis subjects contained a greater diversity of phylogroup 2 *pyrH* genotypes, compared to gingivitis subjects (Mann-Whitney U-test). One *T. denticola pyrH* genotype (pyrH001) was highly prevalent: detected in 10/10 periodontitis and 6/8 gingivitis subjects. Several subjects harboured multiple *T. denticola pyrH* genotypes. Non-metric multidimensional scaling and PERMANOVA tests revealed no significant differences in overall *pyrH* genotype compositions between periodontitis and gingivitis subjects. In conclusion, our results clearly show that subjects with periodontitis and gingivitis commonly harbour highly-diverse oral treponeme communities.

## Introduction

Bacterial taxa belonging to the genus *Treponema* (commonly referred to as treponemes) are the only spirochete taxa known to commonly colonize the human oral cavity. They typically inhabit oxygen-depleted niches, especially dental plaque biofilms found within the gingival sulcus: the shallow crevice of gum tissue that surrounds the base of the tooth **(1, 2)**. Treponemes constitute a relatively small (e.g. <2%) but significant and taxonomically diverse proportion of the healthy human oral microbiome **(3, 4)**. However, it has been known for decades that their proportions are commonly highly elevated within chronically infected subgingival sites found in individuals with periodontal diseases such as gingivitis and periodontitis (see below) **(5-10)**.

There are an estimated 75-100 ‘species-level’ phylotypes of human oral treponeme bacteria, the majority of which have been identified solely by molecular methods **(4, 11, 12)**. Oral treponemes have been phylogenetically classified into 10 ‘oral phylogroups’ (numbered 1-10) **(11, 13)**, and *ca*. 50 Human Microbiome Taxon (HMT) groups **(4, 14)** based on levels of shared 16S ribosomal (rRNA) gene sequence similarity. There are 9 formally classified *Treponema* species of human oral origin (oral treponeme phylogroup in parenthesis): *Treponema medium* (1), *Treponema denticola* (2), *Treponema putidum* (2), *Treponema lecithinolyticum* (4), *Treponema maltophilum* (4), *Treponema amylovorum* (5), *Treponema socranskii* subsp. *socranskii, paredis*, buccale and ‘04’ (6), *Treponema parvum* (7), *Treponema pectinovorum* (8). ‘*Treponema vincentii*’ (phylogroup 1) is commonly reported in the scientific literature, but remains to be formally reported **(15, 16)**. Three other ‘species-level’ phylotypes of ‘*T. medium*-like’ or ‘*T. vincentii* like’ phylogroup 1 treponemes have been identified (named *Treponema* spp. IA, IB and IC) based on results from a multilocus sequence analysis (MLSA) study based on four highly conserved genes: 16S ribosomal RNA (rRNA), *recA* (recombinase A), *pyrH* (uridylate kinase) and *flaA* (flagellar sheath protein) **(17)**.

The term ‘periodontal disease’ encompasses a spectrum of related infectious-inflammatory diseases affecting the tissues that surround and support the teeth **(18)**. Periodontal diseases range from milder forms, i.e. gingivitis, in which the inflammation is limited to the soft-tissue components of the periodontium; to more serious forms, i.e. periodontitis, in which the underlying bone tissues are also affected **(19)**. They highly prevalent in global populations, and severe periodontitis is the major cause of tooth loss in adults **(20)**. The clinical classification of periodontal diseases has recently been revised and updated. Periodontitis is now characterized based on a multidimensional staging (stage I, initial periodontitis; stage II, moderate periodontitis; stage III, severe periodontitis; and stage IV, advanced periodontitis) as well as a grading system (grade A - low risk, grade B - moderate risk and grade C - high risk) **(21, 22)**. Gingivitis is clinically-assessed using percentage of full mouth bleeding on probing (BOP%) scores according to a dichotomized scale: i.e. at ≥10% BOP without attachment loss or radiographic bone loss; or patients with a reduced periodontium who do not have a history of periodontitis or who have previously been successfully treated **(23)**. The reduced periodontium category may experience attachment loss and hence radiographic bone loss.

Periodontal disease has a complex polymicrobial etiology; typified by elevated populations of proteolytic and anaerobic bacterial species in subgingival plaque biofilm communities. There is a large body of evidence linking *Treponema denticola* with severe forms of periodontal disease **(5, 24-30)**. Molecular studies performed over the past 10-20 years have associated several other oral treponeme ‘species’ with periodontal disease. Several studies have implicated species or phylotypes (HMTs) corresponding to *T. medium*, ‘*T. vincentii*’ or closely related oral phylogroup 1 treponeme taxa with the etiology of periodontitis **(10, 29, 31-35)**. However, uncertainties in taxonomy within this cluster of species/phylotypes hinders accurate etiological associations **(11, 12, 15)**.

Here, we analyzed *pyrH* gene sequences encoded by phylogroup 1 and 2 treponeme bacteria present in human subgingival plaque samples collected from subjects with gingivitis (*n*=8) versus subjects with severe forms of periodontitis (*n*=10). Our results showed that both gingivitis and periodontitis subjects harboured diverse communities of phylogroup 1 and 2 treponemes, as indicated by the respective sets of *pyrH* gene sequences identified. Certain phylogenetic lineages of *pyrH* sequences, including one closely related to *T. denticola* ATCC 35405^T^, were highly prevalent within both gingivitis and periodontitis subject groups. Other treponeme *pyrH* gene lineages showed disease-selective distributions. In summary, our results demonstrate that individuals with both milder and more severe forms of periodontal disease may harbor multiple genetic lineages corresponding to the same species or species-level phylotype of oral treponeme.

## Materials and Methods

### Subject Recruitment

A cross-sectional design was employed to compare the diversity of oral treponemes in subgingival plaque from periodontally-diseased sites in subjects presenting with periodontitis and gingivitis. Chinese subject with a minimum of 20 teeth excluding third molars and teeth planned for extraction, who attended the Reception Clinic, Prince Philip Dental Hospital from July 2009 to February 2015 were chosen from the patient pool and were invited to attend a screening session. Smoking history and dental status were extracted from record of first attendance and were confirmed on the screening visit. Trained dentists examined the participants. A calibration exercise on the clinical operations by experienced dentists was carried out prior to the start of the study. The full mouth %BOP scores, probing pocket depth (PPD), and radiographic bone level **(36)** were recorded after written informed consent.

Subjects fulfilling the following criteria were invited to partake in the study. The inclusion criteria were as follows: Periodontitis (P) group: at least two teeth-sites with PPD ≥ 5 mm in two quadrants **(37)** with radiographic evidence of interproximal alveolar bone loss more than 1/3 root length; gingivitis (G) group: 1) PPD ≤ 3mm on an intact periodontium or PPD ≤ 4mm on a reduced periodontium; 2) no radiographic bone loss in any standing tooth; 3) BOP ≥ 10% **(38)**. The exclusion criteria were 1) presence of conditions suggesting a need for antibiotic prophylaxis prior to periodontal examination and invasive dental treatment; 2) history of systemic disease or taking medications known to be associated with periodontal conditions; and 3) history of periodontal treatment except oral hygiene instructions, or antibiotic therapy in the past 6 months. The periodontal conditions of P and G group participants were classified according to current recommendations **(39)**.

Ethical approval was granted by Institutional Review Board of the University of Hong Kong/Hospital Authority Hong Kong Cluster West (UW 15-641), and the study was performed in accordance with the Declaration of Helsinki.

### Sample collection

Prior to sampling, the target clinical sites were isolated with sterile cotton rolls and air-dried. Supra-gingival plaque was removed by sterile universal curettes and subgingival plaque samples were collected using sterilized Gracey curettes from all the diseased sites (PPD ≥ 5 mm) from the P group participants, or from all subgingival sites from the G group participants, respectively. Samples from each subject were pooled into a 2 ml sterile screw cap micro-centrifuge tube containing 1.0 ml of phosphate buffered saline (PBS; 0.01 M phosphate, 137 mM NaCl, 2.7 mM KCl; pH 7.4). Samples were stored in -70°C until processing. After thawing, subgingival plaque samples were washed twice with PBS, and genomic DNA was extracted using the QIAamp DNA mini kit (Qiagen Group, USA) following the manufacturer’s instructions. DNA was eluted in 100 µl of manufacturer’s elution buffer.

### PCR amplification and DNA sequencing of treponeme *pyrH* genes

The *pyrH* gene sequences from phylogroup 1 and 2 treponemes were respectively amplified using previously described primers sets **(17)**. Phylogroup 1: pyrH-I-F (ATGGTACGGGTCTTATCGGTAG) and pyrH-I-R (TTAACCTATCGTTGTGCCTTTAA); phylogroup 2: pyrH-II-F: (ATGGTAACTGTTTTGTCGGT) and pyrH-II-R (TTAGCCGATTACCGTTCCTT). PCR reactions were carried out on a GeneAmp PCR System 9700 (Applied Biosystems) using a ‘touchdown PCR’ approach with GoTaq flexi DNA polymerase (1U; Promega); with conditions as previously described **(17)**. PCR products were gel purified using QIAquick Gel Extraction Kits (Qiagen Group, USA), and ‘TOPO-cloned’ into pCR2.1 TOPO vectors (TOPO TA Cloning Kit; Invitrogen, Life Technologies, CA, USA). Ligation mixtures were electroporated into *E. coli* DH10B, plated onto Luria-Bertani (LB) 1% agar plates supplemented with kanamycin (50 μg/ml) and X-gal (5-bromo-4-chloro-indolyl-β-D-galactopyranoside, 20 μg/ml), then incubated overnight at 37°C. All transformant colonies (*ca*. 26-53 for each plate) were inoculated into fresh LB + kanamycin both (4.0 ml), incubated overnight at 37°C, and plasmid DNA was purified from each sample (QIAprep Spin Miniprep Kits, QIAgen). All plasmid inserts were Sanger sequenced bi-directionally using M13 forward and reverse primers (Beijing Genome Institute (BGI) Hong Kong, Ltd.).

### Bioinformatics/sequence processing and genotypes assignment

The bi-directional Sanger sequences were aligned, primers sequences were trimmed-off using BioEdit v.7.2.3 **(40)**. The combined full-length sequences were quality filtered and manually checked. Multiple sequence alignments of gene sequences were constructed using MAFFT v7.266 **(41)**. Pairwise-distance matrices amongst the aligned DNA sequences were calculated using the ‘*dist*.*seqs*’ function in Mothur **(42, 43)**. The *cluster* function was used to group the sequences into operational taxonomic units (OTUs) or ‘*pyrH* genotypes’ (as indicated in the text) by sequence identities based on the average neighbor algorithm. The *pyrH* genotypes were defined using 97% sequence identity cut-offs, based on results from previous analyses **(17)**. Alpha diversity estimators including genotype richness, diversity, and depth of sample coverage were calculated using the ‘*summary*.*single*’ functions in Mothur.

### Phylogenetic analyses: phylogeny estimation, recombination and selective pressures detection

Maximum Likelihood (ML) estimation of phylogenetic trees were calculated for the *pyrH* gene sequence datasets using the program GARLI (Genetic Algorithm for Rapid Likelihood Inference) **(44)**. Prior to the ML estimation, the best substitution model and gamma rate heterogeneity were determined using the Akaike Information Criterion implemented in jModelTest2 **(45)**. The best ML tree topology with the Maximum Clade Credibility was adopted with clade supports annotated at the branch nodes **(46)** after 1000 bootstrapping replicates. Bayesian posterior probabilities were calculated for the best tree topologies using Mr Bayes v3 **(47)**. Branch nodes with ≥ 60% support value were shown in the phylograms.

### Subject and group-based comparisons of clinical *pyrH* genotype composition

The absolute abundances of the respective clinical *pyrH* genotypes (i.e. number of plasmid clones encoding the corresponding *pyrH* sequence) within the respective participants and subject groups, were visualized as heat-maps using the R packages superheat and ggplot2 v2.2.1 (http://www.r-project.org/). Prevalence was defined as the frequency of clinical *pyrH* genotype being detected within each of the respective subjects. The clinical *pyrH* genotype compositions in each subject were also analyzed using Non-metric MultiDimensional Scaling (nMDS). ML trees containing representative *pyrH* genotypes were used to compute generalized UniFrac distance metrics **(48, 49)**. Results were visualized by 2-dimensional nMDS ordination in R using the phyloseq package **(50)**.

### Statistical tests of demographic and clinical parameters

Statistical analysis was performed using SPSS software 25 (SPSS Inc., Chicago, IL, USA) and R package software (3.5.2). The *p* value threshold was set to 0.05. Descriptive statistics were conducted to analyze the clinical parameters. Assumption tests for normality and equality of variance were performed prior to all statistical analyses. Standard errors were calculated to estimate the sampling errors on age, full mouth BOP scores (%), full mouth PPD ≥ 5 mm (%), number of sites sampled, and mean PPD for sampled sites. The interquartile ranges were calculated to estimate the sampling errors on standing teeth, %BOP scores for sampled sites, and the numbers of cloned *pyrH* sequences and numbers of *pyrH* genotypes. Gender differences between the two clinical groups were compared using Fisher’s Exact test. Independent samples T-test was employed to compare age, full mouth %BOP scores, and the number of sites sampled between the P and G groups. One sample T-test was used to compare the mean PPD values for sampled sites. Mann-Whitney U test was used to compare the numbers of standing teeth, %BOP scores for sampled sites, the numbers of cloned *pyrH* sequences and numbers of *pyrH* genotypes between the P and G groups.

## Data availability

All full-length cloned *pyrH* sequences were deposited in the NCBI GenBank with accession numbers **MT091982-MT093184 (Table S5)**.

## Results

### Clinical evaluation of periodontal status in the two subject groups

Eighteen adult subjects with periodontitis (**P**, *n* = 10) or gingivitis (**G**, *n* = 8) were recruited with informed consent. Their respective demographic profiles and full mouth clinical parameters are summarized in **Table 1** and **Table 2**. Three of the P subjects were classified as having stage IV grade C periodontitis, six P subjects having Stage III grade C periodontitis, and one P subject having Stage III grade B periodontitis, according the recently revised classification system **(Table 2) (20, 21)**. The mean age of P group subjects (35.8 ± 9.6 years) was significantly younger than that of the G group (49.5 ± 5.3 years; *p* = 0.002; Independent samples t-test). Consistent with their disease status, the P group subjects (76.4 ± 16.9%) had significantly higher numbers of sites that bled on gentle probing [reported as full mouth bleeding on probing (BOP) scores] compared to the G group (30.4 ± 18.4%; *p* < 0.0001; Independent samples T-test).

**Table 1.** Summary of demography and clinical parameters within periodontitis (P) and gingivitis (G) subject groups.

|  |  | Periodontal condition |  | Statistical test performed | P-value |
| --- | --- | --- | --- | --- | --- |
|  |  | Periodontitis (P) (n = 10) | Gingivitis (G) (n = 8) |  |  |
| Categories |  |  |  |  |  |
| Age in years | (Mean ± SD) | 35.8 ± 9.6 | 49.5 ± 5.3 | t | 0.002 |
| Gender | male | 7 | 3 | Fisher's Exact | 0.45 |
|  | female | 3 | 5 |  |  |
| Standing teeth | median | 27 | 26 | Mann-Whitney U | 0.633 |
|  | interquartile range | 3 | 2 |  |  |
| Full mouth BOP scores (%) | (Mean ± SD) | 76.4 ± 16.9 | 30.4 ± 18.4 | t | <0.0001 |
|  | range | 63.6-100 | 12.4-60.7 |  |  |
| Full mouth PPD ≥ 5 mm (%) | (Mean ± SD) | 20.8 ± 17.10 | 0 | t | 0.004 |
|  | range | 2.7-52.3 | 0 |  |  |
| Profile of sampled sites |  |  |  |  |  |
| No. of sites sampled | (Mean ± SD) | 32.0 ± 24.6 | 157.8 ± 7.5 | t | <0.0001 |
|  | range | 4-69 | 144-168 |  |  |
| BOP for sampled sites (%) | median | 99.3 | 21.5 | Mann-Whitney U | <0.0001 |
|  | interquartile range | 6.0 | 33.0 |  |  |
| Mean PPD (mm) for sampled sites | (Mean ± SD) | 5.9 ± 0.4 | ND |  |  |
|  | range | 5.3-6.5 | ND |  |  |

*Note:* The pocket probing depth (PPD) values for all G group subjects were ≤ 3 mm, and were not recorded (ND). BOP = bleeding on probing. The p-values < 0.05 are indicated with bold text.

**Table 2.**
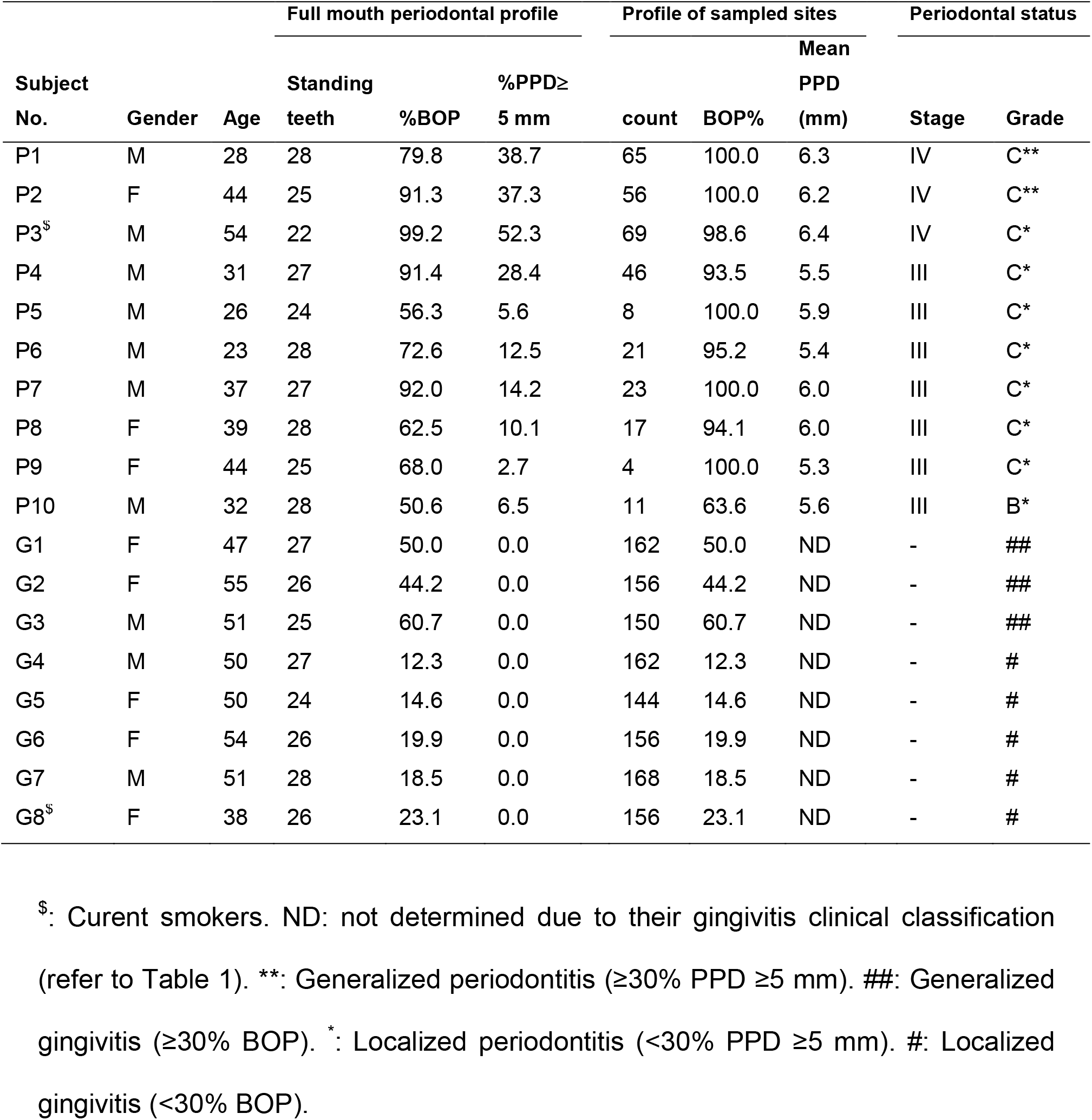
Summary of demography, clinical parameters and periodontal status for each subject.

After clinical examination, a pooled, multi-site sample of subgingival plaque was carefully collected from each subject for nucleic acid purification and molecular analysis. The respective numbers of sites sampled for each subject, as well as the corresponding BOP scores (%) and probing pocket depths (PPD, in mm) for the sampled sites, are summarized in **Tables 1** and **2**. In the P subjects, the median BOP for sampled sites was 99.3%, and the PPD for the sampled sites was 5.9 ± 0.4 mm. This indicated that the vast majority of the sampled sites corresponded to clinically diseased sites. In the G subjects, *ca*. one fifth of the sampled sites tested positive for BOP, indicating that subgingival plaque was collected from a mixture of clinically diseased and non-diseased (clinically asymptomatic) sites.

### Identification of clinical *pyrH* genotypes present within subgingival niches in the two groups

Two sets of PCR primers were used to respectively amplify full-length *pyrH* gene sequences from oral phylogroup 1 and 2 treponemes present in the subgingival plaque samples collected from each subject. These two PCR primer sets have previously been shown to specifically amplify *pyrH* gene sequences from a diverse set of treponeme isolates belonging to oral phylogroup 1 and 2 **(17)**. PCR amplicons of *ca*. 700 bp were successfully obtained using both primer sets for each subject **(Table S1)**. The 36 PCR amplicons were respectively cloned into ‘TOPO’ (pCR2.1) plasmids, to create TOPO plasmid clone libraries of phylogroup 1 and 2 *pyrH* genes for each subject. A subset of the resultant plasmid clones obtained from each of the 36 libraries (range: 26-53) were sequenced bi-directionally. A total of 1,204 quality-filtered, full-length *pyrH* gene sequence reads (heron described as ‘cloned *pyrH* sequences’) were recovered from the clinical cohort **(Table S1)**. A total of 536 cloned *pyrH* sequences were obtained using oral treponeme phylogroup 1 primer set, and 668 cloned *pyrH* sequences were obtained using oral treponeme phylogroup 2 primers. Specificity was 100%, with no ‘off-target’ sequences detected. All of the phylogroup 1 *pyrH* gene amplicons were 687 bp in length (average G+C content 46.0 ± 1.2%), encoding (putative) PyrH proteins of 228 amino acids (aa). All phylogroup 2 *pyrH* gene amplicons were 696 bp in length (average G+C content 40.8 ± 0.7%), encoding PyrH proteins 231 aa in length.

The *pyrH* gene sequences (*n* = 1,204) were clustered into 34 ‘*pyrH* genotypes’, based on a 97% (average neighbour) sequence identity cut-off (**Figure S1** and **Table S2**). This cut-off value was used based on results from a previous study of *pyrH* gene sequence diversity across seven different oral treponeme species/phylotypes **(17)** (see discussion section). These 34 *pyrH* genotypes were named pyrH001−pyrH034, according to the respective total numbers of cloned *pyrH* gene sequences that corresponded to each genotype.

The number of *pyrH* genotypes identified within the P and G subject groups is summarized in **Table 3**. There was no significant difference in the median number of cloned *pyrH* gene sequences respectively obtained from the P and G groups (median = 61.5 for both groups); nor for the number of cloned *pyrH* sequences obtained using the phylogroup 1 or 2 PCR primer sets (*n* = 30 or 31.5 for both groups). There was no significant difference in the median number of phylogroup 1 *pyrH* genotypes identified in the P and G groups (*n* = 3.5 and *n* = 3, respectively). However, the median number of phylogroup 2 *pyrH* genotypes identified in the P group (*n* = 3) was higher than that identified within the G group (n = 1.5; *p* = 0.009; Mann-Whitney U test; see **Table 3**).

**Table 3.** Summary of *pyrH* genotypes and numbers of cloned *pyrH* gene sequences recovered from the periodontitis and gingivitis subject groups.

|  |  | Periodontitis | Gingivitis | *P*-value<sup>a</sup> |
| --- | --- | --- | --- | --- |
| Number of *pyrH* genotypes identified | median<br>range | 7<br>4-12 | 5<br>4-7 | **0.034** |
| Number of cloned *pyrH* gene sequences obtained | median<br>range | 61.5<br>59-81 | 61.5<br>55-83 | 1.000 |
| Number of phylogroup-1 *pyrH* genotypes identified | median<br>range | 3.5<br>2-6 | 3<br>3-4 | 0.696 |
| Number of cloned *pyrH* gene sequences corresponding to phylogroup-1 treponeme species/phylotypes | median<br>range | 30<br>29-30 | 30<br>29-30 | 0.573 |
| Number of phylogroup-2 *pyrH* genotypes identified | median<br>range | 3<br>2-6 | 1.5<br>1-3 | **0.009** |
| Number of cloned *pyrH* gene sequences corresponding to phylogroup-2 treponeme species/phylotypes | median<br>range | 31.5<br>30-52 | 31.5<br>26-53 | 0.829 |

*Note:* <sup>a</sup> Mann-Whitney U test. The *p*-values <0.05 are indicated with bold text.

A total of 326 cloned *pyrH* gene sequences corresponded to the pyrH001 genotype, making it the most frequently detected genotype within the cohort (**Figure 3** and **Table S2**). PyrH001 was the most frequently detected genotype in both the P subjects (*n* = 135) and the G subjects (*n* = 191; discussed further below). Thirteen *pyrH* genotypes (pyrH022−pyrH034) were each represented by a single cloned *pyrH* sequence (**Figure 4**).

The alpha-diversity and richness of the respective *pyrH* genotypes identified within the two clinical groups were determined using the Chao1, ACE, Shannon, Simpson’s and Good’s coverage estimators (summarized in **Tables S3** and **S4**). All individuals were sampled adequately as indicated by Good’s coverage values, which ranged from 0.9 to 1.0 for both the phylogroup-1 and -2 *pyrH* genotypes.

### Taxonomic classification and phylogenetic relationships between *pyrH* genotypes

#### 1. Phylogroup 2 oral treponemes including *T. dentcola* and *T. putidum*

Phylogenetic relationships between the 34 *pyrH* genotypes were inferred by Maximum Likelihood (ML) estimation (**Figures 1** and **2**) with reference to results from a previous study **(17)**. In our analysis, we included *pyrH* gene sequence data from several oral treponeme strains whose genomes have been deposited in the eHOMD database **(14)**. The ML analyses included 22 oral phylogroup-1 and 31 oral phylogroup-2 treponeme strains, respectively. Full-length *pyrH* sequences from *Treponema lecithinolyticum* ATCC 700332 (phylogroup 4), *Treponema maltophilum* ATCC 51939 (phylogroup 4) and *Treponema socranskii* subsp. *socranskii* ATCC 35536 (phylogroup 6) were included as outgroups. The best substitution model used for the ML trees was GTR+G+I, as determined by jModeltest2 **(45)**.

**Figure 1.**
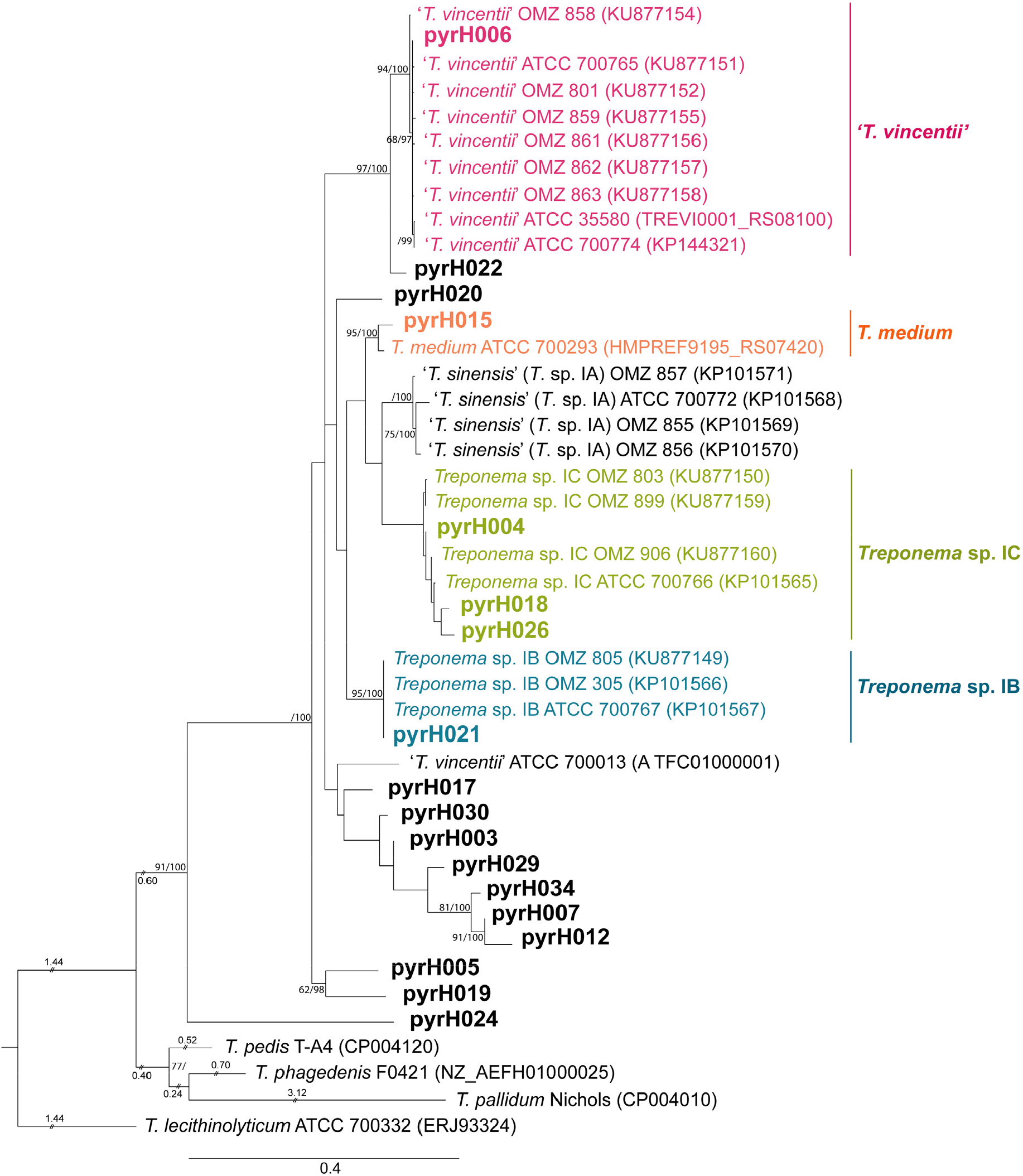
Maximum Likelihood (ML) phylogenetic tree of *pyrH* genes from oral treponeme phylogroup 1 taxa. The maximum clade credibility tree topology is supported by bootstrapping for 1,000 replicates (first number) and Bayesian posterior probabilities (second number) reported as percentage values separated with a forward slash “/” symbol beside the branch nodes. Only values over 60% are shown. Treponeme genotypes identified in this study are indicated in bold type. Excessively long branches have been trimmed in proportion to the scale bar (indicated with “//” on the respective branches). The scale bar indicates 0.4 nucleotide changes per position. The respective oral treponeme species (phylogroups) are indicated with different color shadings as follows: ‘*T. vincentii*’, pink; *T. medium*, orange; *Treponema* sp. IC, lawn-green and *Treponema* sp. IB, cyan. *Treponema pedis* T-A4, *Treponema phagedenis* F0421, *Treponema pallidum* Nichols and *Treponema lecithinolyticum* ATCC 700332 were included as outgroup species.

Eighteen of the 34 clinical *pyrH* genotypes were classified as phylogroup 1 treponeme taxa (**Figure 1**), with 16 corresponding to phylogroup 2 treponemes (**Figure 2**). Twelve of the 34 *pyrH* genotypes could be taxonomically assigned to a previously identified oral treponeme species or phylotype **(17)** (**Table S2**), and are represented with different colours in **Figures 1** and **2**. The other 22 *pyrH* genotypes were classified as corresponding to oral phylogroup 1 treponemes of uncertain taxonomic standing (*Treponema* sp. I*) or to oral phylogroup 2 treponemes of uncertain taxonomic standing (*Treponema* sp. II*), and are shown in black in **Figures 1** and **2**.

**Figure 2.**
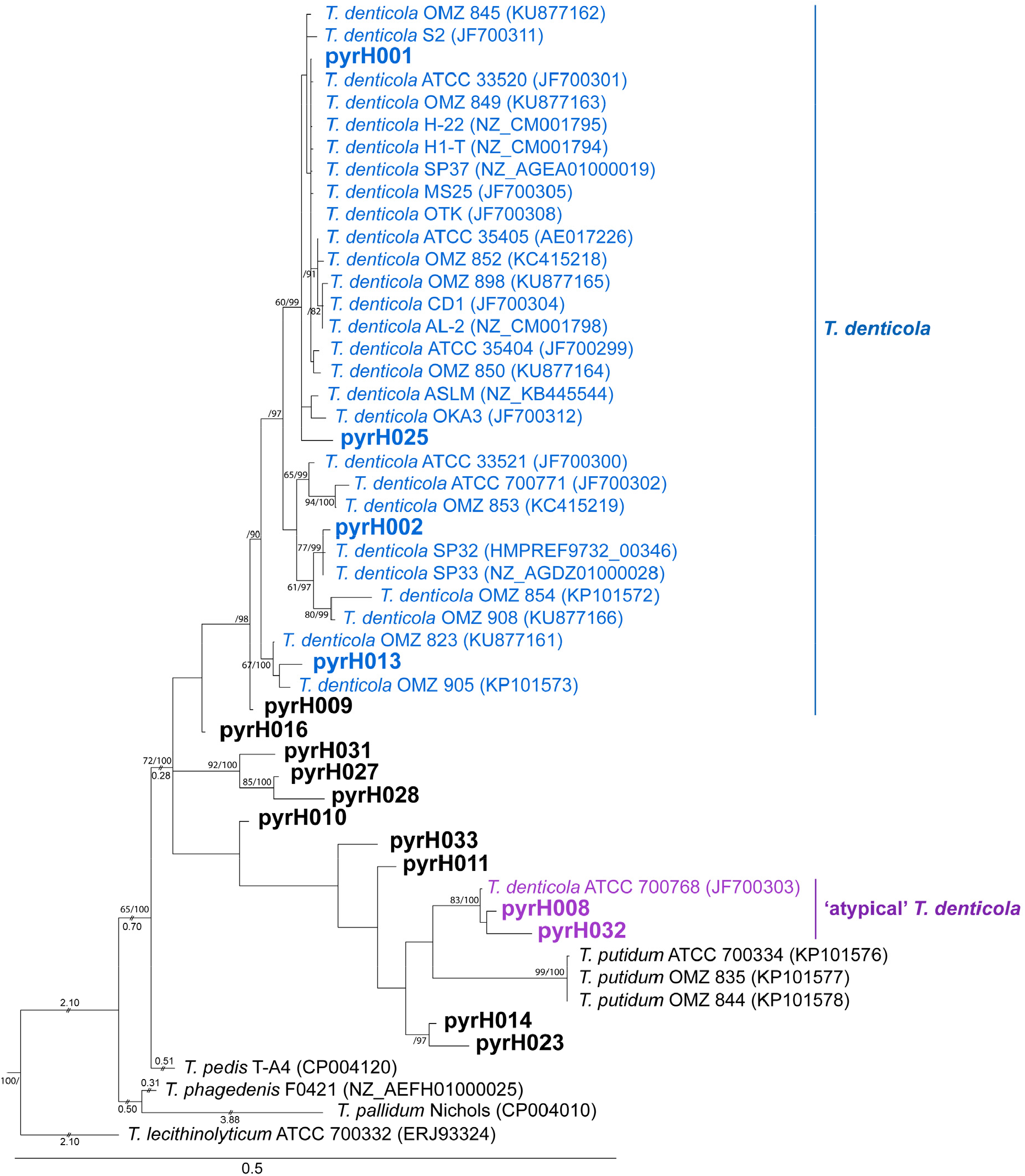
Maximum Likelihood phylogenetic tree of *pyrH* genes from oral treponeme phylogroup 2 taxa. The respective oral treponeme species (phylogroups) are indicated with different color shadings as follows: *T. denticola*, blue and ‘atypical’ *T. denticola* strains, purple. A scale bar indicates 0.5 nucleotide changes per position. For other explanatory details refer to Fig. 1.

Four *pyrH* genotypes (pyrH001, pyrH002, pyrH013 and pyrH025) corresponded to *T. denticola* (coloured blue in **Figure 2**). Genotype pyrH001 was located in a large clade containing the ATCC 35405 type strain (**Figure 2**). Genotype pyrH002 was most closely related to the SP33 and SP32 strains (mean identity 99.3%), whilst pyrH013 formed a distinct clade with the OMZ 823 and OMZ 905 strains (mean identity 97.8%).

Genotypes pyrH008 and pyrH032 formed a distinct clade with the ATCC 700768 (OMZ 830) strain, which was originally classified (and deposited) as an ‘atypical’ strain of *T. denticola* (mean identity 97.6%) **(51)** (shown in purple in **Figure 2**, and discussed further below). The *pyrH* gene sequence of this ‘atypical’ strain is more closely related to those from isolates of *T. putidum* than from the other *T. denticola* strains sequenced to date. Genotypes pyrH027, pyrH028 and pyrH031 formed a distinct clade (within clade mean identity 95%), which was well separated from the other phylogroup 2 taxa. None of the clinical *pyrH* genotypes identified within the cohort corresponded to *T. putidum*, which is the only other formally characterized species in phylogroup 2. Furthermore, we did not detect any *pyrH* gene sequences corresponding to the closely-related (pathogenic, mammalian host-associated) species of treponeme, such as *Treponema phagedenis, Treponema pallidum*, or *Treponema pedis* (**Figure 2**).

#### 2. Phylogroup 1 oral treponemes including *T. medium* and ‘*T. vincentii*’

We identified *pyrH* genotypes corresponding to 4 out of the 5 previously classified species/phylotypes of oral phylogroup 1 treponemes **(17, 52)**, including: ‘*T. vincentii*’ (pyrH006), *T. medium* (pyrH015), ‘*Treponema* sp. IB’ (pyrH021), and ‘*Treponema* sp. IC’ (pyrH004. pyrH018, pyrH026). We did not detect *Treponema* sp. IA (‘*T. sinensis*’) in this subject group. It may be noted that the sequence of the *pyrH* gene from the ATCC 700013 (OMZ 779) strain of ‘*T. vincentii*’ is highly anomalous to the *pyrH* genes encoded by all other known strains in this species **(17)**. We identified seven *pyrH* genotypes that clustered with this strain, including two that were prevalent in the cohort and were detected with high frequency (pyrH003 and pyrH007; see discussion section) (**Figure 3**). The sequence of the pyrH021 genotype was identical to the *pyrH* gene present in all 3 known strains of *Treponema* sp. IB: OMZ 305 (which was formerly classified as ‘*T. vincentii* Ritz A’), OMZ 805 and ATCC 700767 (OMZ 806) **(17)**. The pyrH022 and pyrH020 genotypes represented distinct single taxon lineages. The pyrH005 and pyrH019 genotypes formed a distinct, well-separated clade. The pyrH024 genotype was a notable outlier, and was phylogenetically divergent from all other known *pyrH* gene sequences.

**Figure 3.**
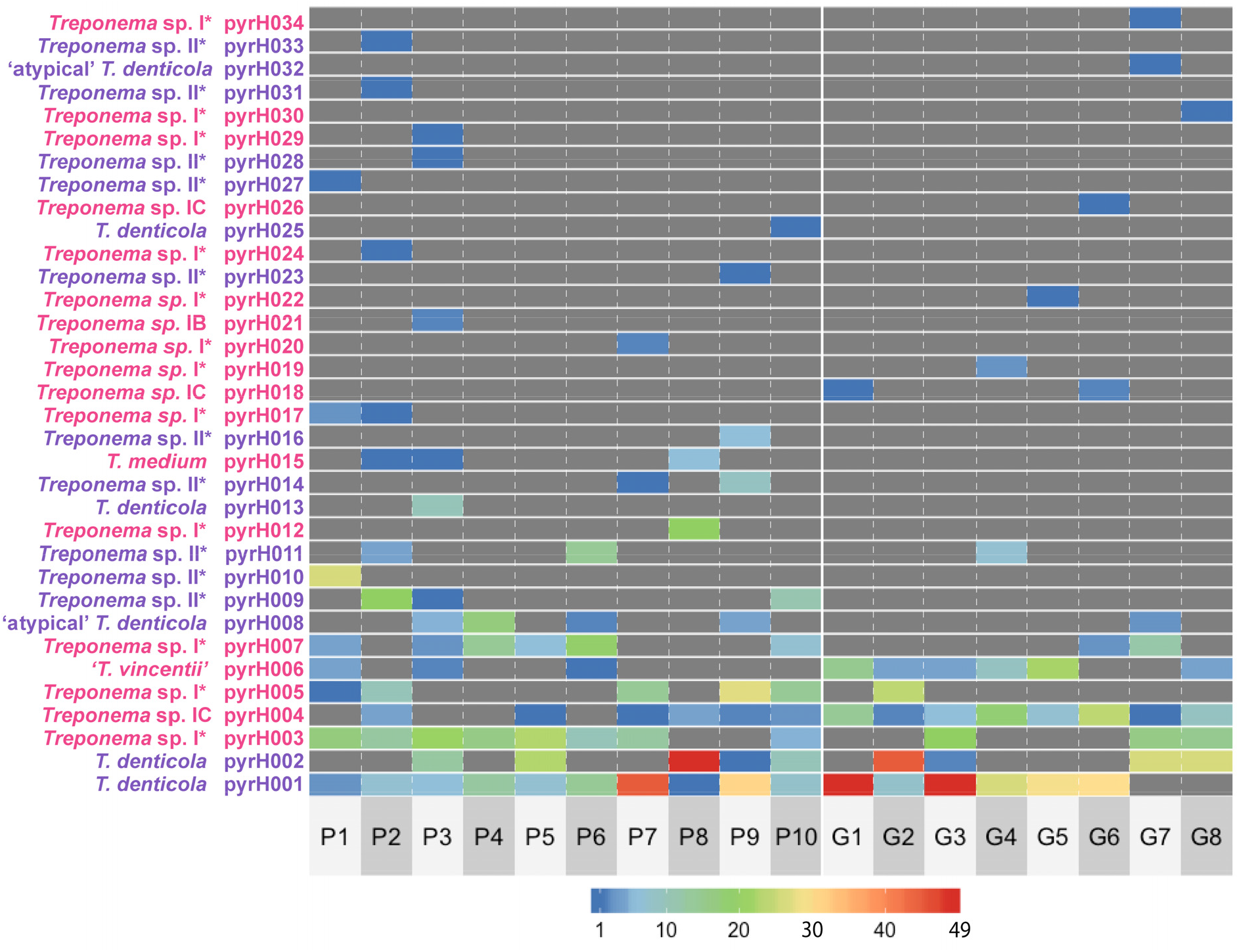
Heat-map showing the clonal abundance of the 34 oral treponeme *pyrH* genotypes detected within each subject. The respective identities of the 34 treponeme *pyrH* genotypes (pyrH001-034) are shown on the y-axis. The identities of the 18 subjects are indicated below the x-axis (P1-10 = periodontitis group; G1-8 = gingivitis group). The respective clonal abundances of each treponeme *pyrH* genotype (i.e. the numbers of cloned *pyrH* gene sequences assigned to each *pyrH* genotype) in each subject are represented in colour according to the scale bar shown below the heat map (zero values are colored dark grey). The pink and purple font colors indicate *pyrH* genotypes belonging to oral phylogroups 1 and 2, respectively.

### Distributions of *pyrH* genotypes in the two clinical groups

The absolute counts of cloned *pyrH* gene sequences within each of the respective subjects were visualized in a heat-map (**Figure 3**). The prevalence of each *pyrH* genotype within the P and G clinical groups (i.e. detection/non-detection in each subject) is shown in a bar chart in **Figure 4**. The *T. denticola* pyrH001 genotype was the most prevalent *pyrH* genotype in the cohort, which was found in 10/10 of periodontitis subjects and 6/8 gingivitis subjects. Although its prevalence was slightly higher in the P group, a higher number of cloned *pyrH* gene sequences were obtained from the G group subjects (*n* = 191 vs. *n* = 135; see **Figure 4**). The *T. denticola* pyrH002 genotype was also prevalent in the cohort, being detected in 5/10 P subjects and 4/8 G subjects, with approximately equal numbers of cloned *pyrH* sequences obtained from P versus G subjects. Genotype pyrH003 (*Treponema* phylogroup I*) was highly prevalent in P group subjects (8/10) and less prevalent in the G subjects (3/8). Conversely, the *Treponema* sp. IC pyrH004 and pyrH006 genotypes were highly prevalent in the G subjects (8/8 and 6/8, respectively), but less prevalent in the P subjects (6/10 and 3/10, respectively).

**Figure 4.**
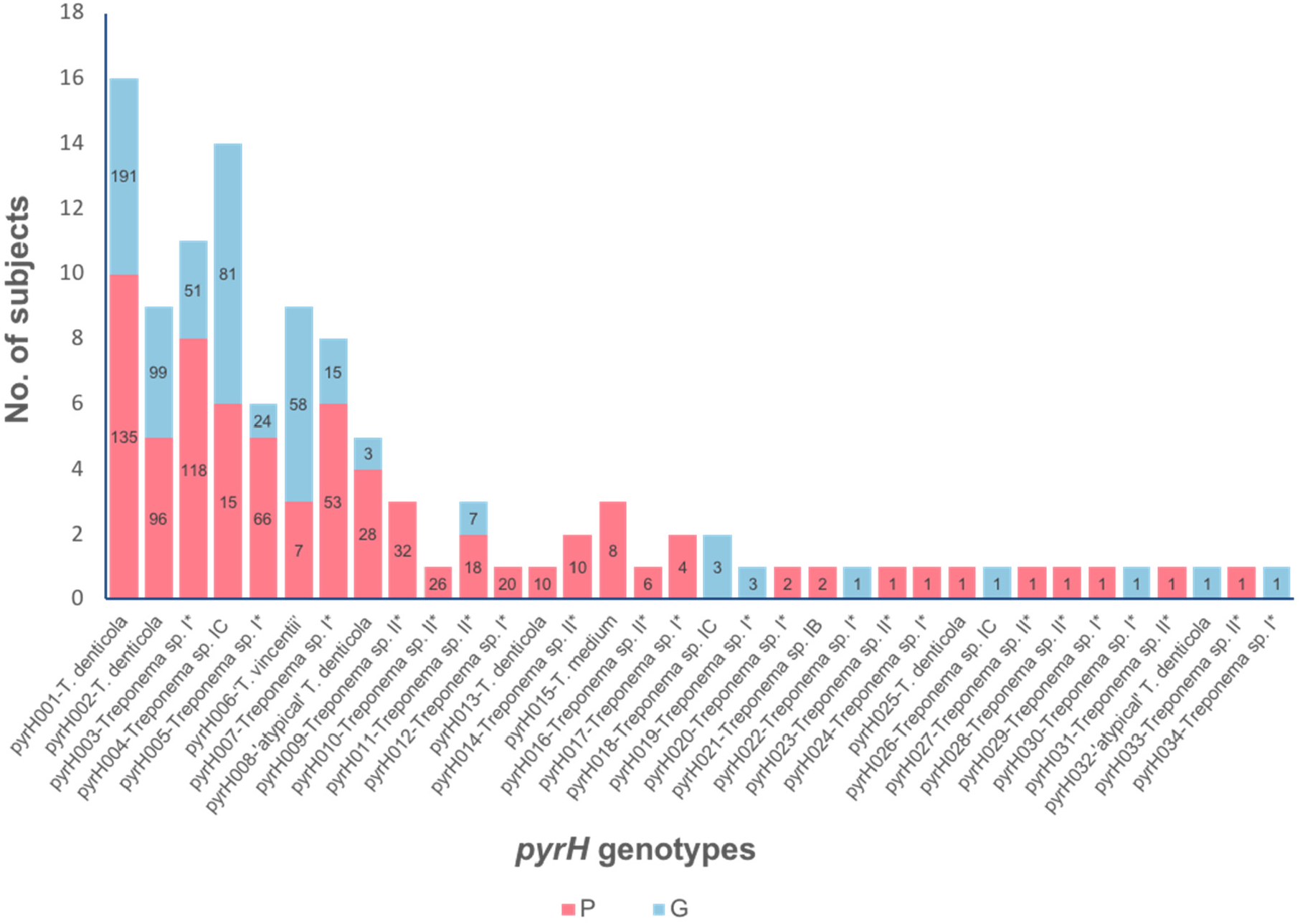
Prevalence of the oral treponeme *pyrH* genotypes detected in the periodontitis (P) and gingivitis (G) subject groups. The heights of the bars indicate the detection frequency of each of the *pyrH* genotypes (pyrH001-pyrH034) in the P and G subject groups (i.e. indicate the total number of subjects that tested positive for each *pyrH* genotype). The red and blue colours represent the prevalence within the P and G subject groups, respectively. The number values in the bars indicate the respective clonal abundances of the 34 *pyrH* genotypes (i.e. corresponding number of cloned *pyrH* gene sequences) in the P and G subject groups (out of a total of 1,224 cloned *pyrH* gene sequences). Thirteen pyrH genotypes (pyrH022-034) were identified by a single cloned *pyrH* gene sequence.

Non-metric Multi-Dimensional Scaling (nMDS) ordination analyses of the clinical *pyrH* phylogroup-1 and -2 genotypes composition based on the generalized UniFrac distance are respectively shown in Panels A and B in **Figure 5**. The segregation between the *pyrH* genotypes from P and G groups are more apparent in the phylogroup 1 data. However, Permanova tests indicated that the differences between the two clinical groups were not significant (Permanova overall R^2^ = 0.099, *p* = 0.121; phylogroup1 R^2^ = 0.157, *p* = 0.018; phylogruop2 R^2^ = 0.114, *p* = 0.097).

**Figure 5.**
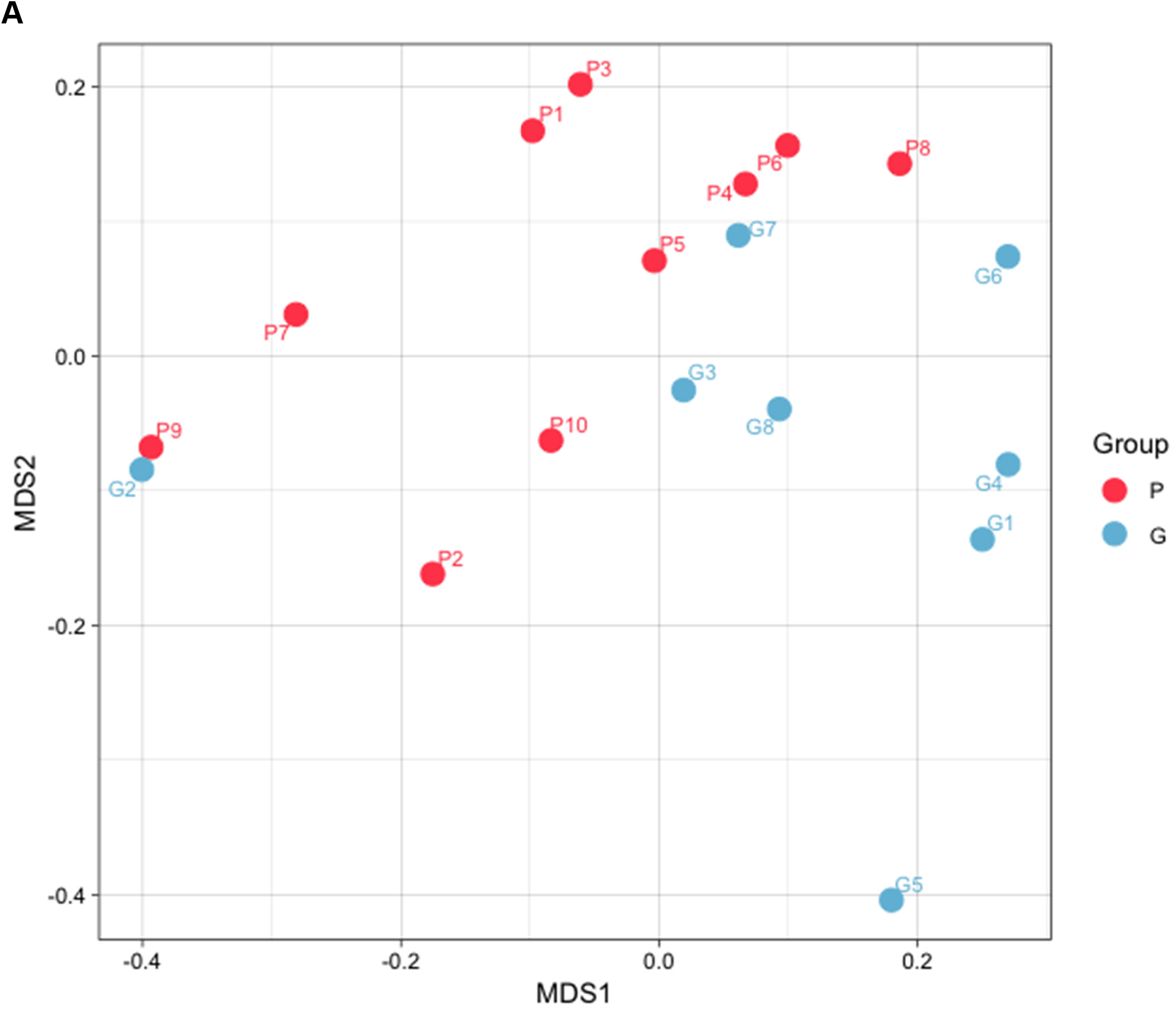
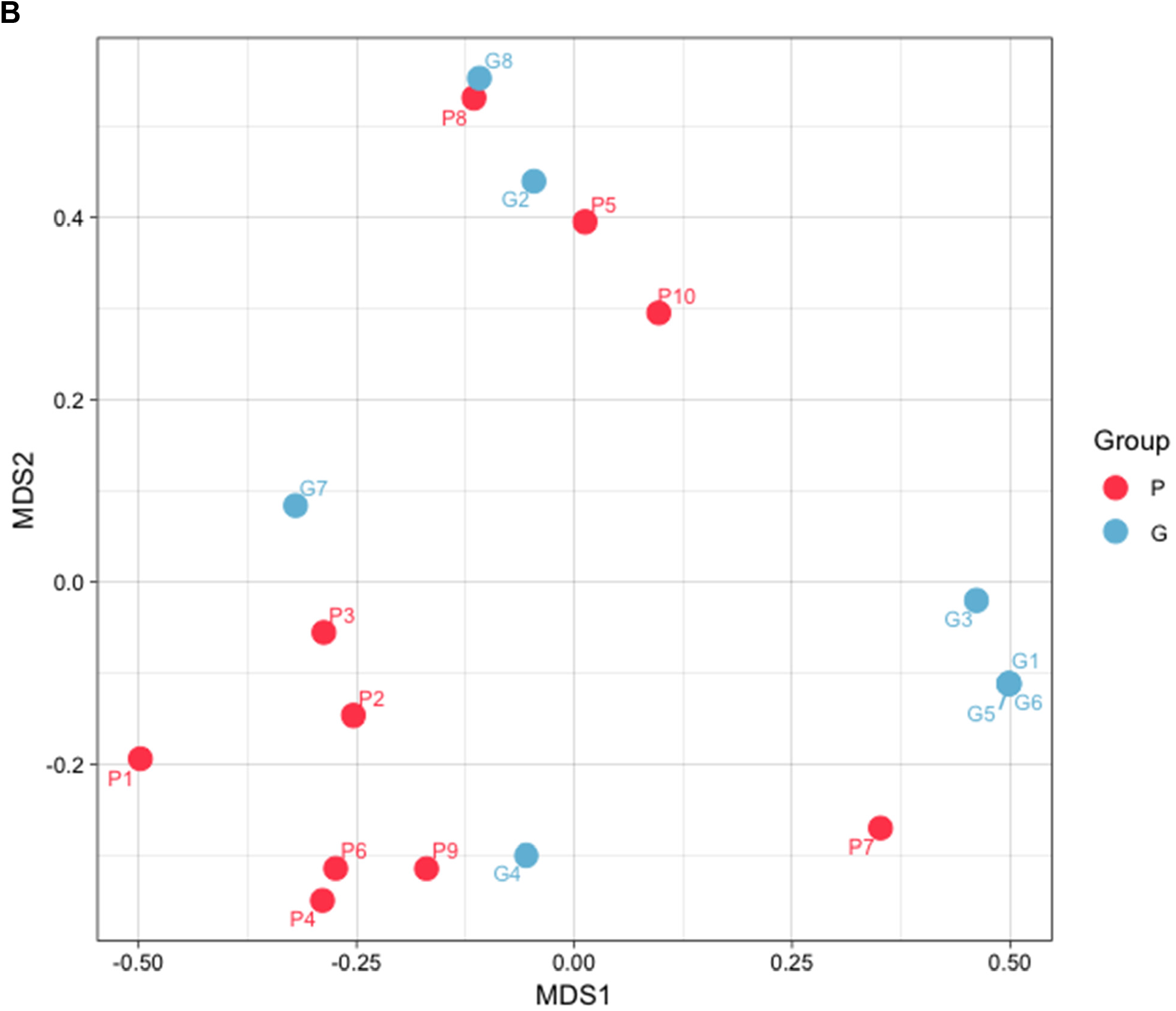
Non-metric multidimensional scaling (nMDS) of oral treponeme *pyrH* genotypes present within each subject. The nMDS was performed using a distance matrix calculated using a generalized UniFrac algorithm based on a single maximum likelihood (ML) tree containing all treponeme *pyrH* genotypes sequences detected (*n*=34). Red solid circles represent the 10 periodontitis subjects (P1-P10); Blue solid circles represent the 8 gingivitis subjects (G1-G8). Panel **(A):** Plot showing the nMDS of treponeme phylogroup-1 genotypes (R^2^ = 0.157, *p* = 0.018). Panel **(B):** Plot showing the nMDS of treponeme phylogroup-2 genotypes (R^2^ = 0.114, *p* = 0.097).

## Discussion

Here, we analyzed differences in treponeme populations within subgingival plaque biofilms sampled from human subjects with gingivitis and periodontitis, using the *pyrH* gene as a diagnostic genetic biomarker. We had two major aims. The first aim was to determine if there were notable differences in the distributions of *T. denticola*, ‘*T. vincentii’, T. medium* and closely-related oral treponeme phylotypes within patients with periodontitis, versus those suffering from gingivitis. The second aim was to establish whether or not subjects with periodontitis or gingivitis harboured multiple genetic lineages (i.e. different clinical ‘strains’) of the same treponeme species/phylotype within their respective oral cavities.

We focused on phylogroup 1 and 2 oral treponemes for the following reasons. Numerous studies have associated *T. denticola* with periodontal disease, and an assortment of virulence factors have been identified for this species **(28, 29, 53-55)**. However, *T. putidum* is phenotypically and phylogenetically very similar to *T. denticola*, yet appears to be relatively rarely detected in clinical cohorts **(10-12, 51, 56, 57)**. In addition, the etiological roles of commonly-detected phylogroup 1 oral treponeme species (phylotypes), such as ‘*T. vincentii*’ and *T. medium* remain to be firmly established **(5, 12, 15, 55, 58)**.

The *pyrH* gene was selected as it has previously been shown to have good strain-resolving abilities as well as a robust correlation with the phylogeny of the 16S rRNA gene within clinical isolates of *T. denticola, T. putidum*, ‘*T. vincentii*’, *T. medium* and other species-level phylotypes within oral treponeme phylogroups 1 and 2 **(17, 59)**. This highly conserved, single-copy ‘housekeeping’ gene has previously been used to assign taxonomy and to infer phylogenetic relationships in MLSA studies conducted within other bacterial species including *Vibrio* spp. **(60)** and *Photobacterium* spp. **(61)**. Our sequencing data shows that the two respective *pyrH* PCR primer sets used had excellent specificity, with no ‘off-target’ hits detected. Due to the limited number of oral treponeme genomes published to date, it is difficult to accurately gauge their clinical detection ‘range’. However, they were previously shown to successfully amplify *pyrH* gene amplicons from all clinical strains tested **(17, 59)**, which validates their effectiveness.

Our results clearly demonstrate that individuals with periodontitis or gingivitis commonly harbor multiple lineages of the same *Treponema* species/phylotype within their respective oral cavities. For example, subjects P3 and P10 contained at least 3 distinct *pyrH* genotypes that corresponded to *T. denticola*. Subject G6 harbored at least 3 different *pyrH* genotypes that corresponded to the *Treponema* sp. IC phylotype **(17, 62)** (**Figure 3**).

The most prevalent and frequently detected *pyrH* genotype (pyrH001) corresponded to a large cluster of previously-identified *T. denticola* isolates that includes the ATCC 33520 and ATCC 35405^T^ strains (**Figure 2**). Strains within this cluster were originally isolated from subjects who resided in various countries across Asia, Europe and North America **(17, 59)**. The second most prevalent *pyrH* genotype (pyrH002) was phylogenetically most closely related to two North American *T. denticola* strains (SP32, SP33) whose genomes were sequenced at the Broad Institute and subsequently deposited in the NCBI GenBank (SP32: GCA_000413095.1; SP33: GCA_000338475.1; unpublished). Taken together, this suggests that there are several *T. denticola* lineages (clinical strains) that are highly prevalent within global populations.

Our results showed that subgingival plaque samples collected from periodontitis subjects contained a greater detectable diversity of phylogroup 1 and 2 oral treponeme *pyrH* genotypes than corresponding samples from gingivitis subjects (i.e. a higher median number of *pyrH* genotypes were detected in P subjects) (**Table 3**). In particular, the diversity of phylogroup 2 oral treponemes were significantly higher in periodontitis subjects, than in gingivitis subjects. This is consistent with results from a previous 16S rRNA gene amplicon-based analysis of oral treponemes in subjects with or without periodontitis **(12)**. In this previous study, periodontitis subjects had significantly higher levels of phylogroup 2 treponeme OTU richness and clonal abundance, with one *T. denticola* lineage (OTU 8P47) having a statistically significant association with periodontitis. Many of the phylogroup 2 *pyrH* genotypes we detected here correspond to as-yet uncharacterized genetic lineages of uncertain taxonomic standing (**Figure 2**). Several of these almost certainly correspond to as-yet uncharacterized lineages of *T. denticola*; however, in the absence of additional genome sequence data, we have been conservative in our taxonomic assignment.

The most frequently detected phylogroup 1 *pyrH* genotypes corresponded to ‘*T. vincentii*’ (pyrH006) and the ‘*Treponema* sp. IC’ phylogroup (pyrH004), which is proposed to constitute a distinct oral *Treponema* ‘species’ **(17)**. The genome sequence of a reference strain of this proposed new ‘species’: *Treponem*a sp. strain OMZ 804 (ATCC 700766) has recently been published **(62)**. It may be noted that the pyrH004 and pyrH006 genotypes were highly prevalent within the gingivitis subjects, and were less prevalent in the periodontitis subjects (**Figures 3 and 4**), but these differences were not statistically significant (*p* = 0.0915 and 0.1534, Fisher’s Exact test). However, it is important to stress that both the periodontitis and gingivitis subjects contained a wide diversity of phylogroup 1 and 2 oral treponeme taxa in their subgingival niches; but had notable differences in their ‘species’ composition.

The diversity of oral treponeme ‘major surface protein (*msp*) genotypes’ has previously been surveyed in a cohort of subjects with or without periodontitis **(59)**. Consistent with the results presented here, the most frequently detected *msp* genotype (NP2_6) corresponded to the *T. denticola* ATCC 35405 type strain, which was equally prevalent in both subject groups. However, due to the very high levels of Msp sequence diversity within oral treponemes, it is not possible to directly correlate many of the *pyrH* genotypes described here with the previously defined *msp* genotypes with high levels of confidence.

Our study has several limitations. Firstly, the sample size of our cohort is relatively small. It is large enough to clearly demonstrate that individuals with periodontitis or gingivitis commonly contain multiple *T. denticola* lineages (as well as those of other treponeme phylotypes). However, it may not be large enough to indicate how many different ‘clinical strains’ of the respective species/phylotypes are typically present in an individual subject’s oral cavity. In addition, as we used pooled multi-site samples, our genetic analysis presents an overview of phylogroup 1 and 2 treponeme distributions within subgingival plaque biofilm niches in the oral cavities of Hong Kong Chinese individuals. It does not indicate if multiple lineages of *T. denticola* or other treponeme species/phylotypes occupy the same clinical site, or reside within distinct sites. It should also be noted that the respective numbers of cloned *pyrH* gene sequences (i.e. clonal abundance) corresponding to each *pyrH* genotype are only semi-quantitative indicators of the actual abundance of oral treponeme cells encoding the respective *pyrH* genes on their chromosomes. Thus, the respective abundances of the various treponeme taxa detected in the different subjects cannot be directly compared. Additional detailed studies are required to answer or further investigate these important questions.

## Conclusions

Subjects with periodontitis and gingivitis harbor a wide diversity of *T. denticola*, ‘*T. vincentii*’ and other oral phylogroup 1 and 2 treponeme taxa within subgingival niches. Our results suggest that there are profound differences in treponeme species/phylotype community composition in subjects with periodontitis versus gingivitis. Notably, individual subjects commonly harbor multiple ‘clinical strains’ of the same oral treponeme species within their respective oral cavities.

## Acknowledgements

This work was financially supported by the Research Grants Council (RGC) of Hong Kong through a General Research Fund (GRF) grant #17105115, awarded to RMW. WKL acknowledges financial support from the Health and Medical Research Fund (HMRF grant #03141636) as well as the University of Hong Kong (Research and Conference Grant, Small Project Fund #09176107). We thank Mr. Raymond Tong and Ms. Joyce Yau of the Central Research Laboratory of the HKU Faculty of Dentistry for their excellent technical support.

